# CCT-217: A Landmark Gene Silencing Therapeutic for Weight Loss That Enhances Lean Mass Composition, Reduces Visceral, Subcutaneous Fat and Restores Metabolic Health in Obesity through Browning

**DOI:** 10.1101/2024.12.30.630715

**Authors:** Raj Reddy, Inderpal Singh

## Abstract

According to the CDC, between 2017 and March 2020, 41.9% of U.S. adults aged 20 and older had obesity, with 9.2% experiencing severe obesity, affecting over 100 million and 22 million adults, respectively. Obesity prevalence increased from 30.5% in 1999-2000 to 41.9% in 2017-2020, while severe obesity rose from 4.7% to 9.2%. As per CDC projection, by 2030, nearly half of the US adults will be obese, with a quarter of them being severely obese. Current therapeutic approaches predominantly focus on appetite suppression and caloric restriction. This approach results in both fat mass and lean mass loss and thus compromises long term metabolic health. Preserving lean mass during weight loss is crucial. Remodeling of the white fat into beige fat through the process called browning offers a promising alternative to appetite suppression dependent weight loss, and complements with reduction of visceral fat, marrow adipose tissue, improve liver steatosis, enhancing insulin sensitivity, reduced cardiovascular risk and suppression of inflammation. Here, we present CCT-217, a groundbreaking siRNA therapy targeting CB1R and ZFP423 genes, with strong adipose tissue-specific tropism with nil brain and liver penetration in diet-induced obese (DIO) C57BL/6J male mice model (n=7 control, n=5 treated). Mice received 7 subcutaneous doses over 20 days (initial half-dose, followed by once after three days CB1R siRNA 2.0 mg/kg and ZFP423 siRNA 1.8 mg/kg dose). CCT-217 induced browning of the WAT and resulted in a significant 26% reduction in body weight with a modest 12% decrease in feed intake after the 4th dose, preserved normal clinical behavior and nutrition. Lean mass composition significantly improved to 79% versus 62% in controls, while total fat mass decreased by 43.5%. Notably, visceral fat depots, including retroperitoneal (59.4%), mesenteric (54.3%), and gonadal (42.0%) fat showed substantial reductions, alongside a reduction of 61.7% inguinal white adipose tissue (iWAT). Histological analysis revealed extensive browning of iWAT, reduced adipocyte size, and increased adipocyte count, indicating enhanced thermogenesis and adipose tissue remodeling. Liver pathology showed significant improvement, with reduced hepatocellular adipose vacuolation and the absence of fibrosis, inflammation, or macrophage infiltration, highlighting the hepatoprotective effects of CCT-217. Metabolic markers demonstrated a 20% reduction in plasma cholesterol and decreased systemic inflammation (21% reduction in CRP), accompanied by significantly improved leptin sensitivity and favorable trends in insulin sensitivity. Mechanistically, CCT-217 achieved 42-fold silencing of CB1R and 14.5-fold silencing of ZFP423 specifically in iWAT, with no off-target effects observed in liver or brain. In conclusion, CCT-217 emerges as a landmark therapy that effectively reduces visceral fat, enhances lean mass composition, and restores metabolic health through precise, tissue-specific gene silencing. These findings position CCT-217 as a transformative and highly promising next generation therapeutic for addressing obesity and its associated complications.

## INTRODUCTION

Obesity is a global health crisis that significantly increases the risk of metabolic disorders, including type 2 diabetes, cardiovascular disease, and Metabolic Dysfunction-Associated Steatotic Liver Disease (MASLD) (Ng et al., 2014; Saklayen, 2018; Younossi et al., 2016). Despite the availability of pharmacological treatments, current therapies often fail to achieve sustained weight loss while preserving lean mass (Christoffersen et al., 2022; Karakasis et al.; Wing & Hill, 2001). Effective strategies addressing both adipose accumulation and energy expenditure are urgently required. The cannabinoid type 1 receptor (CB1R) and the zinc-finger transcription factor ZFP423 are two promising therapeutic targets for obesity due to their roles in complementary metabolic pathways. CB1R overactivation promotes hyperphagia, lipogenesis, and reduced energy expenditure, leading to fat accumulation and metabolic dysfunction (Cota, 2007; Di Marzo et al., 2001). While early CB1R antagonists, such as rimonabant, reduced weight and improved metabolism, adverse central nervous system (CNS) effects resulted in their withdrawal (Després et al., 2005). In contrast, peripherally restricted CB1R inverse agonists, such as CRB-913, JD-5037 and TM-38837 and INV-202 have demonstrated substantial anti-obesity effects, including reductions in visceral fat and improved insulin sensitivity, with mild to moderate CNS-related side effects at higher doses but are limited by partial efficacy (Engeli et al., 2005; Högberg et al., 2024; Morningstar et al., 2023; Nordisk., 2024; Tam et al., 2012).

ZFP423, on the other hand, maintains white adipocyte identity by suppressing thermogenic gene expression (Hepler et al., 2017; Mengle Shao & Rana K. Gupta, 2019). It inhibits uncoupling protein 1 (UCP1) expression, thereby preventing browning of white adipose tissue (WAT) and limiting energy expenditure (Hepler et al., 2017; Mengle Shao & Rana K. Gupta, 2019). ZFP423 suppression triggers the conversion of white adipocytes into thermogenically active beige adipocytes, increasing systemic energy expenditure and protecting against diet-induced obesity (DIO) (Shao et al., 2016). Importantly, even visceral fat depots resistant to browning respond to ZFP423 inhibition (Hepler et al., 2017). CB1R inhibition and ZFP423 suppression target distinct yet synergistic mechanisms. While CB1R silencing reduces energy intake, lipogenesis, and adipose accumulation, ZFP423 silencing enhances energy expenditure through adipose browning (Shao et al., 2016; Zhao et al., 2020). Combining these pathways could overcome limitations of monotherapies by simultaneously reducing fat mass and preserving lean body composition.

We introduce CCT-217, a novel siRNA-based therapy designed to silence CB1R and ZFP423 specifically in adipose tissue. This targeted approach mitigates off-target effects in the liver and brain, ensuring tissue-specific efficacy without adverse events. In 20-days the DIO C57BL/6J male mice with 7 subcutaneous doses injected subcutaneously after every three days, CCT-217 achieved a remarkable 26% reduction in body weight without plateauing, with only an 18% decrease in feed intake compared to the untreated controls. After the 10th day of the study there was only 12% reduction in feed intake compared to the untreated control group. Echo MRI analysis revealed a 43.5% reduction in total fat mass, while lean mass composition increased significantly from 62% to 79% in the CCT-217 treated DIO mice. Notably, visceral adipose depots, including retroperitoneal (59.4%), mesenteric (54.3%), and gonadal (42.0%) fat, exhibited substantial reductions, addressing a critical driver of metabolic complications.

Histological analysis of inguinal WAT demonstrated significant browning, characterized by increased multilocular adipocytes, smaller cell size, and higher adipocyte density, indicative of enhanced thermogenic activity. Liver histopathology revealed reduced hepatocellular vacuolation without signs of fibrosis, inflammation, or macrophage infiltration, highlighting the hepatoprotective effects of CCT-217. CCT-217 also improved systemic metabolic parameters, including a 20% reduction in plasma cholesterol, decreased systemic inflammation (21% reduction in CRP with undetectable IL-6), and restored leptin sensitivity. Insulin sensitivity showed favorable trends, further supporting improved metabolic homeostasis. Mechanistically, CCT-217 achieved robust and tissue-specific gene silencing, with 42-fold reduction in CB1R and 14.5-fold reduction in ZFP423 expression in inguinal WAT.

Compared to existing therapies, such as GLP-1/GIP agonists, peripherally restricted CB1R antagonists, and browning agents, CCT-217 offers superior outcomes by integrating the benefits of reduced adipose accumulation and enhanced energy expenditure. Unlike GLP-1/GIP agonists, which are associated with lean mass loss, CCT-217 preserves and improves lean body composition while significantly reducing visceral adiposity (Karakasis et al.). Its targeted action minimizes CNS and liver-related off-target effects, addressing limitations of traditional CB1R inverse agonists.

CCT-217 represents a transformative advancement in obesity therapy. By combining CB1R inhibition and ZFP423 silencing, it effectively reduces both visceral and subcutaneous fat mass, induces WAT remodeling into beige adipose tissue, enhances energy expenditure, preserves lean body composition, and improves metabolic health with a mild reduction in feed intake suggesting its strength in protecting from the nutrition deplition. These findings position CCT-217 as a promising candidate for clinical translation, offering a novel, tissue-specific approach to combat obesity and its associated metabolic disorders.

## Materials and Methods

### Study Design, administration and Monitoring

This study evaluated the therapeutic efficacy of CCT-217 (a siRNA formulation targeting CB1R and ZFP423) in a **diet-induced obesity (DIO) mouse model**. The experiment utilized 12 male **C57BL/6J DIO mice** aged 17-18 weeks, sourced from the Jackson Laboratory. The mice were individually housed in sterile, ventilated cages under controlled conditions: temperature of 19-25 °C, relative humidity of 30-70%, a 12-hour light/dark cycle, and a minimum of 15 fresh air changes per hour. They were fed a high-fat rodent diet (60 kcal%, Research Diets, D12492) and provided with purified water ad libitum.

After a two-week acclimatization period, mice were randomized into two groups based on body weight: **Vehicle Control Group (n=7)** received Dulbecco’s phosphate-buffered saline (DPBS). The **CCT-217 Treatment Group (n=5)** was administered with CCT-217 siRNA ZFP423 at 1.75 mg/kg and CB1R siRNA at 2 mg/kg, both siRNA mixtures were encapsulated in SM-102 lipid nanoparticles. The siRNA formulation was delivered subcutaneously, once every 4 days and a total 7 doses were injected during the length of the study, over the study period of 20 days.

During acclimatization, baseline measurements of body weight and food intake were recorded. Post-dosing, animals were observed for clinical signs for one hour. Body weight, food intake, and rectal temperature were recorded daily.

### Biochemical and Hormonal Analysis

At study termination, blood was collected via intracardiac puncture. Plasma was separated by centrifugation at 3500 g for 5 minutes at 2-8 °C and analyzed for Liver enzymes (ALT, AST, ALP), Lipid profile (total cholesterol and triglycerides), Hormonal markers (TSH, T3, T4) and Inflammatory markers (CRP, IL-6). ELISA kits (R&D Systems, Invitrogen) were used following manufacturer protocols. This approach builds on published findings regarding adipose tissue inflammation and browning (Bellitto et al., 2024; N. Ottaway et al., 2015; Shen et al., 2014).

### Gene Expression

RNA was extracted from adipose tissue using the RNeasy Mini Kit (Qiagen). Tissue homogenization was performed in TRI reagent under ice-cold conditions. Complementary DNA (cDNA) synthesis utilized the High-Capacity cDNA Reverse Transcription Kit (Thermo Fisher). GAPDH and 18S rRNA were used as reference genes in quantitative PCR (qPCR) assays to normalize expression of CB1R and ZFP423 genes. Primers and probes were run on the ABI Prism 7500 system, with relative gene expression calculated using the ΔΔCT method. Previous studies validated the relevance of these target genes in metabolic regulation and adipogenesis (Gupta et al., 2010; M. Shao & R. K. Gupta, 2019).

### Body Composition Analysis: EchoMRI

Body composition was analyzed using EchoMRI-500™ (EchoMRI, USA), a non-invasive quantitative magnetic resonance imaging system. Mice were transported to the EchoMRI facility in CCMB Hyderabad. Measurements of lean body mass, fat mass, and water content were obtained while ensuring minimal stress to the animals. The data provided precise insights into body fat distribution and composition changes due to CCT-217 treatment.

### Histopathology

Adipose and pancreatic tissues were fixed in 10% neutral buffered formalin, processed, and embedded in paraffin. Sections of 3-5 µm were stained with hematoxylin and eosin (H&E) for histological assessment. Adipocyte area, number, and pathological changes were evaluated under light microscopy. The methodology draws upon approaches used in metabolic studies (Hao et al., 2015; Nickki Ottaway et al., 2015).

### Statistical Analysis

Data were analyzed using ANOVA and Student’s t-tests with a significance threshold of p<0.05. Results were expressed as mean ± standard deviation (SD). Percent changes in metabolic parameters were calculated using normalized data, consistent with analyses in studies (Han et al., 2019; Li et al., 2008).

### Ethical Approvals and Compliance

This study was conducted in accordance with established ethical guidelines for animal research and approved by the Institutional Animal Ethics Committee (IAEC). The study adhered to the regulations outlined by the **Committee for Control and Supervision of Experiments on Animals (CCSEA)** under Animal License No. 2173/PO/RcBiBt/S/2022/CCSEA, dated 27th June 2022. The protocol for the study was reviewed and approved by the IAEC under Protocol No.: GVSAP/IAEC/2024-035. The study followed guidelines to minimize pain, discomfort, and stress to the animals.

### Animal Welfare Measures

Animals were monitored twice daily for clinical signs. Specific interventions were planned for animals in anticipation of any distress during treatments. Humane endpoints were established to minimize suffering, including early euthanasia in cases of severe distress or body weight loss exceeding acceptable limits.

All animals in the control and CCT-217 treated groups remained clinically healthy during the whole of the study and at the end, euthanasia was performed via an overdose of isoflurane, ensuring a painless and humane end. Tissue samples were collected post-mortem for further analysis. Animal housing and enrichment were designed to provide a comfortable and stimulating environment, following CCSEA and IAEC recommendations.

## RESULTS

### Significant Weight Loss with Minimal Reduction in Food Intake Following CCT-217 Therapy

CCT-217 treatment in diet-induced obese (DIO) mice induced a progressive and substantial reduction in body weight with only mild reductions in cumulative feed intake, demonstrating its efficacy and metabolic advantage. By Day 20, CCT-217-treated mice achieved a 26% adjusted weight loss compared to non-diabetic DIO control mice **(Fig. 1A)**. The weight reduction was consistent and pronounced, with an early decline of ∼10% observed by Day 5, indicating the rapid onset of therapeutic effects. Notably, the dosing regimen involved only 7 subcutaneous doses administered once every 4 days over the 20-day period, emphasizing the durability and efficacy of CCT-217 despite infrequent dosing.

**Figure 1:**
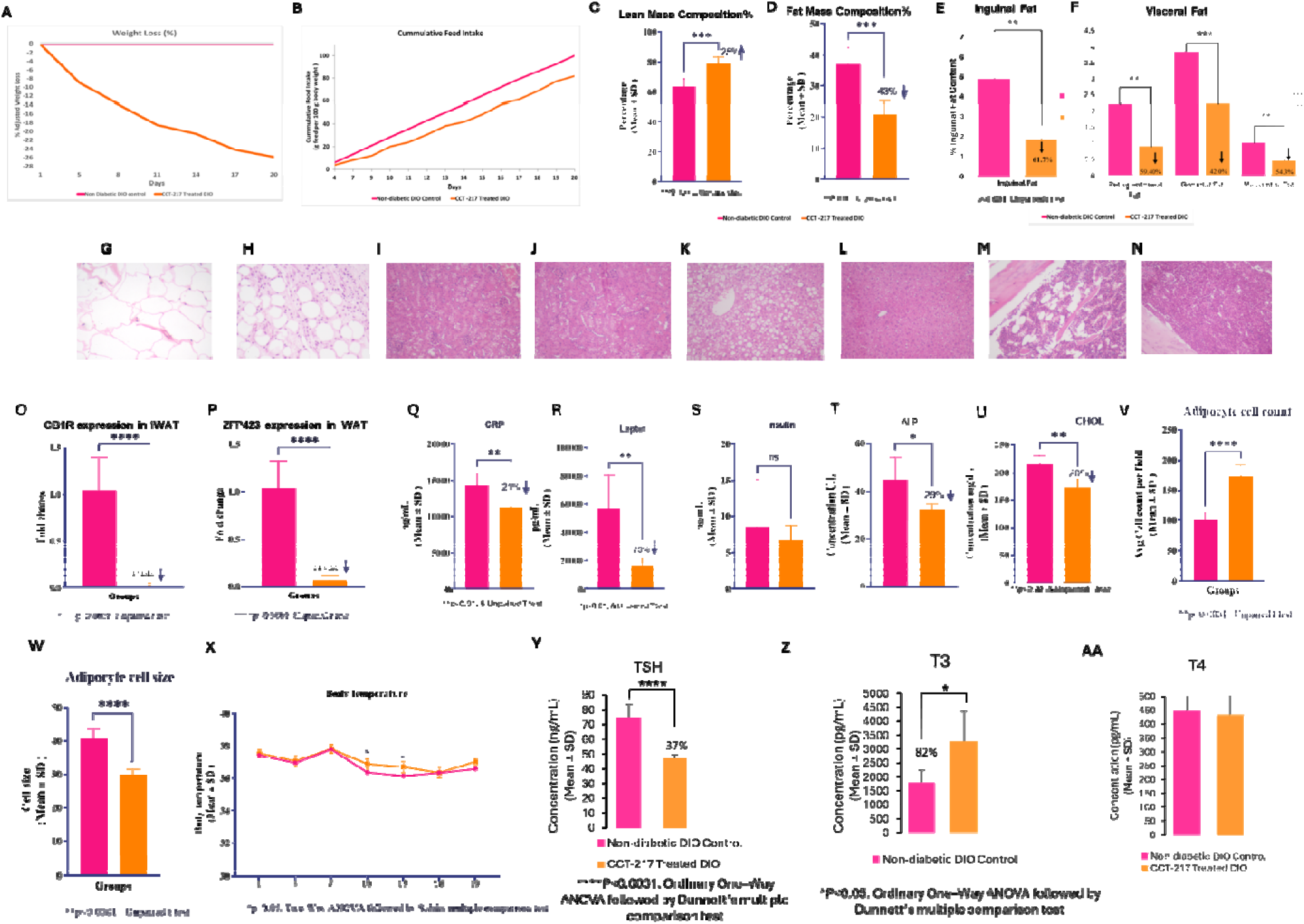
Effects of CCT-417 treatment in non-diabetic DIO mice across metabolic, histopathological, biochemical, and inflammatory parameters are compared to control mice. A significant 26% weight loss in CCT-217 treated mice group was observed vs control **(A)**, along with a mild reduction in cumulative feed intake **(B)**. Echo MRI results revealed lean mass composition improved by 25 % **(C)** and fat mass reduced by 43% in the treated mice **(D)**. Significant reduction in inguinal fat i.e. 61.7% **(E),** retroperitoneal (59.4%), gonadal (43%) and mesenteric (54.3%) fat pads were also observed in the CCT-217 treated group **(F)**. H&E-stained sections of inguinal white adipose tissue (iWAT) from non-diabetic DIO Control **(G)** and CCT-417 Treated **(H)** mice. Control iWAT shows enlarged, hypertrophic adipocytes, while treated iWAT exhibits smaller, more uniformly sized adipocytes, indicative of adipose tissue remodeling and reduced hypertrophy following CCT-417 treatment indicates browning. H&E-stained kidney sections from non-diabetic DIO Control **(I)** and CCT-417 Treated DIO **(J)** groups. Control kidneys show signs of lipid accumulation and cellular hypertrophy, while treated kidneys exhibit preserved architecture with reduced lipid deposits and improved histological features, indicating a protective effect of the treatment. Panel **K** represents the liver section of non-diabetic DIO Control mice, showing significant lipid accumulation and signs of hepatic steatosis. Panel **L** is the liver section of CCT-417 Treated DIO mice, demonstrating a marked reduction in lipid accumulation and improved liver histology, indicative of reduced hepatic steatosis and improved metabolic health. Bone marrow H&E-stained sections from Control **(M)** and Treated **(N)** groups reveal a higher prevalence of adipocytes, reduced hematopoietic cellularity in the control group. In contrast, the treated group **(N)** demonstrates improved cellularity, increased myeloid lineage activity, and reduced adipocyte infiltration, suggesting mitigation of osteopenic features. The RT-qPCR results for **CB1R (O)** and **ZFP423 (P)** expression demonstrate significant downregulation i.e., 42 fold and 14.5 fold respectively in the CCT-217 Treated DIO group compared to the non-diabetic DIO Control group. Significant reduction in inflammation is evident through decrease in CRP levels i.e., 21% in treated mice **(Q)** and leptin sensitivity was restored as evident from 70% decrease in its levels in CCT217 treated group **(R)** (*p < 0.01* for both CRP and leptin). Improvement trend in insulin sensitivity is observed **(S).** 29 % reduction in ALP levels (*p < 0.05*) in the CCT-217 treated group indicate improvement in liver function **(T)**. Cholesterol levels were reduced by 20% (*p < 0.01)* in the treated group indicating improvement in lipid profile **(U)**. In the iWAT microscopic analysis, the adipocyte count was increased whereas their size was decreased in the CCT-217 treated group indicating the browning **(V-W).** Rectal temperature measurement did not report fever or hyperthermia in the treated mice **(X).** Significant reduction in thyroid-stimulating hormone (TSH) levels in CCT-417 Treated DIO mice compared to vehicle control, indicating improved thyroid axis feedback regulation (*p < 0.0001*) **(Y)**. Significant increase in triiodothyronine (T3) levels in CCT-417 Treated DIO mice compared to vehicle control, reflecting enhanced T4-to-T3 conversion and increased metabolic activity (*p < 0.05*) an improvement in obesity **(Z)**. Comparable thyroxine (T4) levels between CCT-417 Treated DIO and vehicle control groups, suggesting stable thyroid hormone production **(AA)**.

While assessing the cumulative feed intake we observed a mild reduction of 18% in feed intake in CCT-217-treated mice relative to the controls **(Fig. 1B)**. It is noteworthy to mention that after the 4th dose on day 10 the cumulative feed intake further improved in the treated mice and reached up to 88% of the untreated control DIO mice thus during this period there was only 12% drop in feed. Overall, the food intake increased progressively in both groups. Importantly, the observed weight loss was disproportionate to the modest reduction in caloric intake, suggesting that CCT-217 likely enhances energy expenditure through the thermogenic processes known as browning/beiging to drive substantial fat and weight loss. These results underscore the ability of CCT-217 to achieve clinically significant weight loss with a favorable dosing schedule and without drastic reductions in food intake, positioning it as a promising and metabolically favorable therapeutic for obesity management.

### Fat mass reduction DIO Mice

CCT-217 treatment induced substantial improvements in body composition, favorably shifting the ratio between lean and fat mass. Echo MRI analysis on Day 22 revealed a 25% improvement in lean mass composition (79% in treated vs. 62% in DIO control group, *p*<0.001), demonstrating CCT-217’s ability to preserve and enhance lean tissue mass—critical for maintaining metabolic health during weight loss (Fig. 1c). Concurrently, fat mass composition decreased by 43% (22% in treated vs. 39% in DIO control group, *p*<0.001), underscoring its potent anti-adiposity effects **(Fig. 1D)**. Expert pathological dissection further confirmed a significant reduction in total fat mass. Specifically, inguinal fat decreased by 61.67%, retroperitoneal fat by 59.41%, gonadal fat by 42.01%, and mesenteric fat by 54.35%. Reductions in visceral fat, particularly retroperitoneal, gonadal, and mesenteric depots, are especially noteworthy due to their strong correlation with metabolic disorders and cardiovascular risks **(Fig. 1E-F)**.

CCT-217’s unique ability to reduce fat mass while enhancing and preserving lean mass proportion achieves levels comparable to healthy, chow-fed mice—setting it apart from therapies like GLP-1/GIP agonists, which, while effective at fat reduction, are often limited by concurrent lean mass loss. The targeted depletion of visceral fat combined with muscle preservation highlights CCT-217’s therapeutic potential to improve metabolic health and address obesity-related comorbidities, positioning it as a promising next-generation obesity treatment.

### Comprehensive Microscopic Pathological Evaluation Demonstrated Safety and Efficacy of CCT-217 in Vital Organs and Adipose Tissues Liver

In the CCT-217-treated group, microscopic analysis revealed a clear improvement in liver pathology compared to the vehicle control group. Among the 5 mice treated, 1 mouse (20%) showed no abnormalities at all, whereas none of the control mice exhibited a normal liver. Vacuolation (hepatocellular, midzonal, diffuse) findings were notably improved in the treated group, with 0 cases of moderate or marked/severe vacuolation, compared to the control group where 2 mice (28.6%) displayed moderate vacuolation and 3 mice (42.9%) had marked/severe vacuolation. While mild vacuolation was observed in 4 treated mice (80%), it remained non-progressive and was less severe compared to the control group. The absence of severe pathology and the presence of one completely healthy liver in the treated group underscore the beneficial effects of CCT-217 on liver health.

### Kidney

Kidney pathology in the CCT-217-treated group demonstrated consistent or improved findings compared to the control group. Minimal vacuolation (tubular, cortex, diffuse) was observed in 1 treated mouse (20%), which was lower than the 3 mice (42.9%) in the control group. Mild vacuolation was consistent between groups, affecting 3 treated mice (60%) and an equal number in the control group. Importantly, no progression to severe vacuolation was observed in the treated group. Regarding inflammation (mononuclear cell, pelvis, focal/multifocal), minimal inflammation was seen in 4 treated mice (80%), compared to 3 mice (42.9%) in the control group; however, no cases of mild inflammation were detected in the treated group, whereas 1 control mouse (14.3%) exhibited mild inflammation. Additionally, hydronephrosis (pelvis dilation, unilateral), graded as moderate in 1 control mouse (14.3%), was completely absent in the CCT-217-treated group. These findings suggest that CCT-217 treatment does not exacerbate kidney pathology and provides slight improvements in inflammation and hydronephrosis outcomes.

### Pancreas

Microscopic evaluation of the pancreas revealed no abnormalities in the treated or control groups.

### Bone Marrow

Increased cellularity, specifically of the myeloid type in a diffuse pattern, was observed in the bone marrow of CCT-217-treated mice. Minimal and Mild increases in myeloid cellularity were detected in 2 and 3 out of 5 mice treated respectively, whereas no such findings were present in the vehicle control group. The observed changes suggest a mild activation of hematopoietic activity, which is an adaptive response to systemic metabolic alterations induced by CCT-217. Importantly, the increase in cellularity was limited to a mild grade, with no evidence of pathological progression or adverse bone marrow remodeling, confirming that CCT-217 treatment does not compromise bone marrow integrity or function.

### Brain

Microscopic evaluation of brain tissue revealed no abnormalities in the CCT-217-treated group or the vehicle control group. The absence of pathologically abnormal findings demonstrates that CCT-217 treatment does not adversely impact the central nervous system. This is particularly significant given the importance of ensuring that anti-obesity therapies do not induce neurotoxicity or compromise brain tissue structure and function.

### Thymus

The thymus of CCT-217-treated mice displayed no detectable abnormalities when compared to the vehicle control group. The preservation of thymic tissue integrity indicates that CCT-217 did not negatively affect this critical immune organ. The absence of inflammation, cellular changes, or structural damage highlighted the safety of CCT-217 with respect to immune organ pathology, further supporting its favorable safety profile.

### Inguinal White Adipose Tissue (iWAT)

Analysis of iWAT revealed a reduction in adipocyte size in the CCT-217-treated group, consistent with the treatment’s potent anti-adiposity effects. The reduction in adipocyte size was diffuse in nature, reflecting adipose tissue remodeling and improved metabolic activity induced by CCT-217. These changes align with the observed reductions in overall fat mass and specific fat depots, including visceral fat, as previously reported. The observed remodeling of iWAT highlights CCT-217’s ability to target adipose tissue, promoting favorable metabolic outcomes while maintaining tissue integrity.

The microscopic findings across the pancreas, bone marrow, brain, thymus, and iWAT collectively confirm the safety and efficacy of CCT-217 treatment. Bone marrow showed mild, adaptive hematopoietic activation without adverse findings. Brain and thymus tissues remained entirely unaffected, demonstrating the treatment’s safety for vital systems. Finally, reductions in iWAT adipocyte size reflect the metabolic benefits of CCT-217, further supporting its role in improving body composition while preserving tissue health. These observations provide strong evidence for the therapeutic potential of CCT-217 as a safe and effective treatment for obesity and associated metabolic disorders.

### H&E Stain Results for iWAT Tissue Slice

Histological evaluation of inguinal white adipose tissue (iWAT) stained with Hematoxylin and Eosin (H&E) revealed distinct differences between control and CCT-217-treated groups. In the control group, adipocytes appeared large, unilocular, and relatively uniform in size, with well-defined cell membranes and flattened nuclei pushed to the periphery. The extracellular matrix was sparse, and minimal stromal or inflammatory cell presence was observed, reflecting a metabolically inactive state typical of untreated adipose tissue with high lipid content **(Fig. 1G)**. In contrast, iWAT from CCT-217-treated mice showed significant changes indicative of adipose tissue remodeling and browning **(Fig. 1H)**. Adipocytes were notably smaller and more heterogeneous in size, suggesting lipid mobilization and reduction in fat storage which are observed in the previous studies with CB1R and ZFP423 inhibition (Shao et al., 2016; Yang et al., 2024). Increased stromal cellularity, evidenced by a greater presence of purple-stained nuclei in the interstitial space, indicates ongoing tissue remodeling. Additionally, while a slight increase in extracellular matrix was observed, there were no signs of fibrosis or pathological tissue damage. Overall, these findings highlight the metabolic activation and favorable remodeling of iWAT following CCT-217 treatment, aligning with the observed reductions in fat mass through browning and energy expenditure improved adipose tissue health.

### H&E Stain Results for Kidney Tissue

Histological analysis of kidney tissues stained with Hematoxylin and Eosin (H&E) revealed comparable findings between the vehicle control and CCT-217-treated groups. Both groups exhibited largely intact tubular architecture and well-preserved glomerular structures, with only mild and localized vacuolation observed in tubular epithelial cells. Scattered regions of slight tubular degeneration and minimal cellularity were noted in the control group but did not significantly differ from the treated group **(Fig. 1I-J)**. Overall, the results indicate that CCT-217 treatment does not exacerbate kidney pathology, maintaining renal tissue integrity comparable to the vehicle control group.

### Liver Histological Evaluation: Impact of CCT-217 on Hepatic Steatosis in Diet-Induced Obese Mice

After 20 days of treatment with 7 doses of CCT-217 (administered once every 4 days), histological analysis of liver tissues stained with Hematoxylin and Eosin (H&E) revealed striking improvements in hepatic morphology. Notably, treatment commenced when the mice were 19-21 weeks old, at which point they had already accumulated significant hepatic fat after prolonged exposure to a high-fat diet (HFD). Mice were sacrificed at 23-25 weeks of age, a timeframe sufficient to establish advanced hepatic steatosis, closely resembling MASLD observed in diet-induced obesity. In the vehicle control group, liver sections showed pronounced macrovesicular steatosis, characterized by large lipid droplets filling and displacing the hepatocyte cytoplasm, disrupting normal liver architecture **(Fig. 1K)**. These findings reflect severe fat accumulation caused by prolonged HFD exposure and impaired lipid metabolism.

In contrast, the CCT-217-treated group exhibited a substantial reduction in hepatic fat content. Hepatocytes displayed minimal vacuolation, with intact and uniform cytoplasm, indicating effective mobilization and clearance of hepatic lipids **(Fig. 1L)**. Notably, these improvements occurred with only a minimal 18% reduction in feed intake, underscoring that the observed effects were not simply a result of caloric restriction but of the metabolic actions of CCT-217. This outcome is particularly remarkable considering the chronic HFD exposure and advanced fatty liver at the time of treatment initiation. The histological improvements, reduction in steatosis and preservation of hepatocyte integrity demonstrate that CCT-217 effectively reverses hepatic fat accumulation, offering significant therapeutic potential for improving liver health in diet-induced obesity.

### Bone Marrow Regeneration and Reduced Adiposity Following CCT-217 Treatment in Diet-Induced Obese Mice

Histological analysis of bone marrow sections stained with Hematoxylin and Eosin (H&E) revealed clear differences between the vehicle control group and the CCT-217-treated group **(Fig. 1M-N)**. In the vehicle control group, bone marrow displayed lower cellular density with prominent fat vacuoles occupying significant space within the marrow cavity **(Fig. 1M)**. This observation reflects marrow adiposity, a well-known consequence of prolonged high-fat diet (HFD) feeding. Excessive fat accumulation in the bone marrow is often associated with reduced hematopoietic function, impairing normal blood cell production and signaling a shift toward adipogenesis. In contrast, the CCT-217-treated group exhibited a marked increase in cellularity, particularly within the myeloid lineage, indicating improved hematopoietic activity. The marrow space appeared densely populated with hematopoietic cells, while fat vacuoles were significantly reduced compared to the control group **(Fig. 1N)**. This reversal of adiposity highlights the treatment’s ability to restore bone marrow homeostasis and promote a shift from adipogenesis back to hematopoiesis. The findings demonstrate that CCT-217 treatment effectively counteracts HFD-induced bone marrow adiposity while enhancing hematopoietic regeneration. The observed increase in cellular density and reduction in fat vacuoles suggest a metabolically favorable shift that supports bone marrow function and aligns with the systemic metabolic improvements seen in CCT-217-treated mice. These results underscore CCT-217’s potential in mitigating obesity-related bone marrow dysfunction.

### CCT-217 Treatment Reduces CB1R and ZFP423 Expression in iWAT

Quantitative RT-qPCR analysis revealed a significant reduction in the expression of CB1R and ZFP423 in inguinal white adipose tissue (iWAT) of CCT-217-treated mice compared to diet-induced obese (DIO) control mice. Expression levels were normalized to 18S ribosomal RNA, used as the control housekeeping gene, to ensure accurate quantification. CB1R expression was elevated in the DIO control group but showed a 42-fold reduction following CCT-217 treatment (p < 0.0001), indicating strong suppression of this receptor, which is associated with adipogenesis and lipogenesis **(Fig. 1O)**. Similarly, ZFP423, a transcription factor critical for preadipocyte differentiation into white adipocytes, exhibited a significant 14.5-fold reduction (p < 0.0001) in the treated group compared to controls **(Fig. 1P)**. This pronounced downregulation of CB1R and ZFP423 reflects CCT-217’s robust inhibitory effects on key molecular pathways driving adipose tissue expansion and lipid storage. These findings align with the observed reduction in fat mass and improved adipose tissue remodeling, emphasizing the therapeutic potential of CCT-217 in targeting obesity-associated adipogenic signaling. In a pilot study, we interestingly observed that treatment with ZFP423 siRNA (total 2 mg/kg body weight per week, administered subcutaneously near iWAT as equal halves in two doses over 7 days) resulted in a 26.20-fold decrease in CB1R gene expression, corresponding to 96.18% silencing, thus demonstrating the silencing effect of ZFP-423 on CB1R expression. Similarly, in a separate study (data not shown), we observed that CB1R silencing also led to silencing of the ZFP423 gene in the mice by 3.5-fold or 71.43%. To the best of our knowledge this is first reported evidence for such a feedback loop, where silencing ZFP423 gene potentiates the silencing of CB1R and vice-versa for the browning, thermogenic activation and adipose tissue remodeling. In the pilot study we also observed that subcutaneous delivery of the CCT-217 had absolutely no effect on brain expression of the CB1R thus setting this therapy as safe to avoid any psychiatric side effects.

### CCT-217 Treatment improved Inflammatory and Metabolic Profiles Through Improved Insulin and Leptin Sensitivity in DIO Mice

Quantitative analysis of cytokines and metabolic markers revealed significant improvements in systemic inflammation and metabolic regulation following CCT-217 treatment in diet-induced obese (DIO) mice. These findings provide evidence of positive physiological changes associated with the reduction of adiposity and the restoration of metabolic homeostasis.

**C-Reactive Protein (CRP),** a marker of systemic inflammation, was significantly elevated in the DIO control group, consistent with chronic low-grade inflammation commonly observed in obesity. CRP production is often triggered by adipose tissue-derived inflammatory cytokines, reflecting the overall inflammatory burden in obese subjects. In the CCT-217-treated group, CRP levels were reduced by 21% (p < 0.01), indicating a clear anti-inflammatory effect **(Fig. 1Q).** This reduction is likely a consequence of decreased adipose tissue mass, particularly visceral fat, which is known to release pro-inflammatory mediators such as interleukin-6 (IL-6). By reducing fat depots, CCT-217 diminishes the inflammatory load, thereby lowering CRP levels.

**Leptin,** a hormone primarily secreted by adipose tissue, was significantly reduced by 70% (p < 0.01) in CCT-217-treated mice **(Fig. 1R)**. In obesity, elevated leptin levels reflect both increased adipose mass and leptin resistance, where the brain becomes insensitive to leptin’s satiety and energy-regulating signals. The substantial reduction in leptin following treatment aligns with the significant weight loss and fat mass reduction observed in the treated mice. This decline suggests an improvement in leptin sensitivity, where leptin secretion has normalized in response to the reduced adiposity. Prior studies have demonstrated that weight loss leads to a proportional decline in leptin levels, reflecting improved energy balance and adipose tissue function. CCT-217’s ability to enhance adipose tissue remodeling and reduce fat depots likely restored leptin signaling, contributing to overall metabolic improvement. Conversely, the Leptin management through anti leptin antibodies in the DIO mice have also been known to improve obesity (Zhao et al., 2020).

**Insulin levels** exhibited a trend toward reduction in the CCT-217-treated group, indicating a potential improvement in insulin sensitivity **(Fig. 1S)**. Obesity-induced insulin resistance arises from chronic inflammation and excess lipid accumulation in peripheral tissues, impairing insulin signaling. The reduction in adiposity, systemic inflammation, and improved adipose tissue function seen with CCT-217 treatment would contribute to enhanced insulin sensitivity. As adipose tissue inflammation diminishes and lipid mobilization increases, insulin resistance is alleviated, leading to a normalization of circulating insulin levels. Though not statistically significant in this study, the observed trend suggests early indications of improved metabolic efficiency that may become more pronounced with prolonged treatment.

Overall, the observed reductions in CRP and leptin, along with trends in insulin levels, reflect a systemic improvement in inflammation, insulin sensitivity, and leptin sensitivity following CCT-217 treatment. These results are consistent with known mechanisms linking weight loss, fat mass reduction, and restored adipose tissue function to improved metabolic health. The combination of reduced systemic inflammation and enhanced hormone sensitivity underscores the multifaceted therapeutic potential of CCT-217 in reversing obesity-related metabolic dysfunction.

### CCT-217 Treatment Improves Liver Function and Reduces Cholesterol Levels in Diet-Induced Obese Mice

In this study, clinical chemistry analysis was performed to evaluate liver function, lipid metabolism, and renal indicators in CCT-217-treated diet-induced obese (DIO) mice compared to the vehicle control group. Key liver enzymes, including alanine aminotransferase (ALT), aspartate aminotransferase (AST), and alkaline phosphatase (ALP), were measured to assess hepatic function. A decreasing trend in the ALT and AST levels were observed between the treated and control groups, ALP levels were significantly reduced by 29% (p<0.05) in the CCT-217-treated mice **(Fig. 1T)**. ALP is often elevated in obesity-related hepatic steatosis and inflammation, and its reduction indicates a positive effect of CCT-217 in alleviating liver stress and improving overall hepatic function. Lipid profile analysis revealed a substantial decrease in cholesterol levels in CCT-217-treated mice. The total cholesterol (CHOL) concentration was significantly reduced by 20% (p<0.01) compared to the DIO control group **(Fig. 1U)**. This reduction is particularly relevant as elevated cholesterol is closely associated with metabolic dysregulation in obesity. Overall, the results indicate that CCT-217 treatment significantly improves liver function, as evidenced by the reduction in ALP levels, and promotes favorable changes in lipid metabolism, particularly cholesterol reduction, in diet-induced obese mice. These findings support the therapeutic potential of CCT-217 in alleviating obesity-related hepatic dysfunction and lipid abnormalities while maintaining overall metabolic balance.

### Body Temperature Analysis in CCT-217-Treated Mice

Rectal body temperature measurements were conducted to assess any potential hyperthermic effects of CCT-217 treatment in diet-induced obese (DIO) mice **(Fig. 1X)**. Over the 20-day study period, body temperature remained within the normal physiological range of 36°C-38°C for both the CCT-217-treated and control groups. On Day 10 and Day 13, a modest yet statistically significant increase in body temperature (*p < 0.05*) was observed in the CCT-217-treated group compared to the DIO control group. However, these values were well within the normal range, indicating that the observed elevations were neither excessive nor clinically relevant. By Day 19, body temperature levels in the treated group returned to baseline values comparable to the control group. These results conclusively demonstrate that CCT-217 treatment does not induce clinical hyperthermia at all. The transient elevations in temperature on Days 10 and 13 are likely reflective of increased metabolic activity, potentially due to enhanced energy expenditure and adipose tissue remodeling, which are consistent with the physiological effects of CCT-217. Importantly, the absence of hyperthermia further supports the safety and tolerability of CCT-217 as a therapeutic intervention in obesity.

### CCT-217 Enhances Thyroid Function in DIO Mice

The effects of CCT-217 on thyroid function were evaluated in non-diabetic DIO mice, with untreated controls as the baseline. Mice treated with CCT-217 exhibited a significant reduction in TSH levels (p < 0.0001) compared to controls, indicating improved thyroid axis regulation **(Fig. 1Y)**. Elevated TSH is a hallmark of obesity-associated hypothyroidism, often linked to metabolic dysfunction (Bassett et al., 2010). Additionally, T3 levels were significantly increased (p < 0.05), suggesting enhanced conversion of T4 to T3 **(Fig. 1Z)**, a process critical for energy expenditure and metabolic rate regulation (Silvestri et al., 2018; Roth et. al., 2024). No significant differences in T4 levels were observed, highlighting the specific upregulation of deiodinase activity **(Fig. 1AA)**. Histopathological analysis revealed a smaller but healthy thymus in treated mice compared to untreated controls, indicating reduced systemic inflammation commonly associated with obesity (Hotamisligil, 2006). This suggests improved immune function without compromising thymic health with CCT-217 treatment, supporting its potential as an anti-obesity therapy with systemic metabolic benefits (Yau et al., 2014; Gabay et al., 2010).

## Discussion

A crucial advancement in obesity therapeutics is the development of CCT-217, a novel synergistic siRNA therapy targeting CB1R and ZFP423, which achieves precise subcutaneous adipose tissue specificity without any suppression in the levels of CB1R in the brain or liver. CCT-217’s remarkable subcutaneous adipose tissue-specific tropism posits it with significant therapeutic advantage because it achieves a substantial 26 percent weight loss in just 20 days. This is an exceptional qualitative weight loss because the treated mice had a tremendous improvement in their lean mass composition, comparable to that of healthy lean mice as observed in literature (Nilsson et al., 2016). Notably this was achieved with comparatively very low and less frequent dosing (i.e. once every 3 days @ total 3.75 mg/kg body weight dose) compared to other CB1R targeted therapies or other energy expenditure enhancing. We emphasize that CCT-217’s weight loss represents a significant improvement over existing and investigational GLP/GIPr agonists, as the latter induce primarily quantitative weight loss including substantial lean body mass reduction as reported in some studies to be as high as 20% accompanied by excessive reductions in feed intake, leading to nutritional depletion.

On the contrary, CCT-217 promotes adipose tissue remodeling through browning and improves systemic metabolic health. This CCT-217 treatment induced weight loss resulted in mild reduction in cumulative feed intake of ∼18% compared to ∼45% in case of Tirzepatide (Coskun et al., 2018). Our study concludes that CCT-217 improves the feed intake behavior of the mice along with energy expenditure and browning, rather than over dependence on anorexic mechanisms as seen for GIP/GLPr agonists. Interestingly, after the 4^th^ dose of CCT-217 in the treated mice (i.e., after day 10 onwards) the difference in the feed intake was only 12% compared to the control group’s cumulative feeding which emphasizes the browning of the WAT and energy expenditure in the adipose tissue as the major driver of this weight loss instead of hypo-phagy This dual CB1R and ZFP423 targeted approach provides a compelling mechanistic rationale for superior therapeutic outcomes in obesity, particularly compared to existing mono-therapies like GLP-1/GIP agonists, which primarily reduce fat mass but fail to preserve lean mass or enhance energy expenditure (Karakasis et al.).

Mechanistically, CB1R is a key regulator within the endocannabinoid system (ECS) and plays an important role in energy homeostasis by controlling appetite, lipogenesis, and energy expenditure. Its overactivation in obesity leads to hyperphagia, increased fat accumulation, and systemic inflammation. First generation CB1R targeted inhibitors induced weight loss but were also associated with psychiatric issues, while the next generation peripherally restricted CB1R inverse agonists have shown promising results in reducing visceral adiposity and improving insulin sensitivity, their clinical application remains limited by partial efficacy (Di Marzo, 2008; Silvestri & Di Marzo, 2013). Meanwhile, ZFP423, a zinc-finger transcription factor, is a potent repressor of thermogenic gene expression by inhibiting EBF2 and suppressing UCP1 (Rajakumari et al., 2013; Shao et al., 2016; Shao et al., 2021; W. Wang et al., 2014). By maintaining the white adipocyte phenotype, ZFP423 restricts adipose plasticity and browning potential. Knockout studies demonstrate that silencing ZFP423 can allow for the epigenetic de-repression of thermogenic programs, facilitating the activation of beige adipocytes within white adipose tissue (WAT) depots and enhancing systemic energy expenditure (Shao et al., 2016). Its suppression allows for the reprogramming of white adipocytes into thermogenically active beige cells, even in depots resistant to browning (Shao et al., 2016). This mechanism complements the action of CB1R silencing, which reduces adiposity and systemic inflammation. By targeting both CB1R and ZFP423, CCT-217 leverages a synergistic effect to promote adipose tissue browning, enhance systemic energy expenditure, and achieve superior weight loss outcomes compared to monotherapies, underscoring its transformative therapeutic potential in obesity.

Our findings reveal an unexpected crosstalk between CB1R and ZFP423, where silencing one gene reciprocally downregulated the other. Specifically, CB1R inhibition reduced ZFP423 expression, while ZFP423 silencing further suppressed CB1R. This bidirectional relationship likely reflects shared regulatory pathways governing lipid metabolism, inflammatory signaling, and adipose tissue plasticity. The combined silencing of CB1R and ZFP423 resulted in profound adipose remodeling, including significant fat loss in visceral depots—retroperitoneal (59.4%), mesenteric (54.3%), and gonadal (42.0%)—as well as robust browning of inguinal WAT (iWAT), evidenced by smaller, multilocular adipocytes with thermogenic characteristics. These findings align with the known role of ZFP423 suppression in facilitating adipocyte browning and systemic energy dissipation (Shao et al., 2016).

In comparison to published results, CRB-913, a cannabinoid receptor 1 (CB1R) inverse agonist, administered orally at a total dose of 5 mg/kg body weight daily for 18 days (90 mg cumulative dose), induced a 16.1% weight loss in diet-induced obesity (DIO) mice (Morningstar et al., 2023). Similarly, semaglutide, a GLP-1 receptor agonist, administered subcutaneously at 30 nmol/kg every third day (Q3D) for 18 days, achieved a 19.5% reduction in body weight (Morningstar et al., 2023). Additionally, in a phase 2a trial of monlunabant, a novel oral CB1R inverse agonist, a 10 mg daily dose achieved a 7.1 kg weight loss over 16 weeks, with limited additional weight loss observed at higher doses. However, monlunabant was associated with dose-dependent, mild to moderate neuropsychiatric side effects, including anxiety, irritability, and sleep disturbances (Nordisk., 2024). Importantly, our CCT-217 achieved a superior 26% weight reduction over 20 days with a less frequent dosing regimen of 3.75 mg/kg Q3D and as demonstrated through RT-qPCR validation, no detectable silencing of CB1R was observed in the brain and liver tissue, highlighting its adipose tissue-specific tropism. This superior performance underscores the advantage of CCT-217’s dual-targeting mechanism, which simultaneously promotes adipose tissue browning, enhances thermogenesis, and reduces adiposity (Morningstar et al., 2023).

Notably, the observed weight reduction with CCT-217 was achieved with only a modest 12% reduction in feed intake from dose four onward, highlighting increased energy expenditure rather than caloric restriction as the primary driver of weight loss. This outcome underscores the thermogenic activation of beige adipocytes as a significant mechanism underlying CCT-217’s efficacy. In addition to adipose-specific effects, systemic improvements were evident across multiple metabolic organs. Hepatic pathology improved significantly, with reduced hepatocellular vacuolation and the absence of fibrosis or inflammation, suggesting hepatoprotective effects mediated by decreased lipo-toxicity and systemic inflammation. Bone marrow analyses showed adaptive myeloid activation without adverse remodeling, affirming the safety of CCT-217 therapy.

The observed systemic improvements conferred by CCT-217 likely stem from the secretion of batokines, bioactive molecules released by brown and beige adipocytes during adipose tissue browning. Key batokines could include FGF-21, NRG-4, IL-6, GDF-15, Adiponectin, SLIT2-C, METRNL, and Myostatin, each playing distinct roles in enhancing systemic metabolism. FGF-21 improves hepatic lipid metabolism and leptin sensitivity (Fisher & Maratos-Flier, 2016; Lee et al., 2014), while NRG-4 reduces hepatic lipogenesis and visceral fat mass (G.-X. Wang et al., 2014). IL-6 supports glucose homeostasis and mitochondrial function (Stanford et al., 2013), and GDF-15 reduces inflammation and fosters weight loss (Tsai et al., 2013). Adiponectin and SLIT2-C enhance WAT browning, energy expenditure, and insulin sensitivity (Hui et al., 2015; Svensson et al., 2016). METRNL regulates immune responses and promotes thermogenesis (Rao et al., 2014). Additionally, CB1R signaling has been shown to promote an inflammatory macrophage M1 phenotype, leading to chronic low-grade inflammation, a hallmark of obesity and metabolic disorders (Tian et al., 2017). Its silencing has been shown to alleviate systemic inflammation. The detailed mechanistic insights on biochemical pathways and regulatory networks involved in thermogenesis and energy expenditure in brown and beige adipocytes are explicitly explained in a review written by Chouchani, Kazak and Spiegelman (Chouchani et al., 2019). A large human-based study conducted by Paul Cohen’s group at The Rockefeller University has established a strong statistical link between brown adipose tissue (BAT) activity and improved cardiometabolic health. By analyzing over 134,000 PET/CT scans from more than 52,000 individuals, the study demonstrated that the presence of BAT is independently associated with lower prevalence rates of type 2 diabetes, dyslipidemia, coronary artery disease, and hypertension. This association was particularly pronounced in individuals with obesity, where BAT activity appeared to mitigate the detrimental metabolic effects of excess adiposity. These findings highlight browning’s potential as a therapeutic target to address obesity-related disorders and enhance cardiometabolic and metabolic health (Becher et al., 2021). The combined effects of CCT-217 contribute to the superior metabolic outcomes by addressing key drivers of obesity and its related complications. Our CCT-217 therapy demonstrates a multifaceted mechanism of action by targeting distinct but interconnected pathways in obesity pathophysiology. By silencing CB1R and ZFP423, CCT-217 reduces fat mass, facilitates WAT browning, enhances energy expenditure, and improves systemic inflammation and organ health. This dual-targeting strategy uniquely positions CCT-217 as a transformative therapy for obesity, surpassing the limitations of existing mono therapies by preserving lean mass, reducing visceral adiposity, and improving metabolic and inflammatory markers. Furthermore, CCT-217 has been improved further to achieve its therapeutic effects with an ultra-low dosing regimen of just twice annually, offering compliance for long-term obesity management. These findings highlight the therapeutic potential of CCT-217 in addressing the complex and multifactorial nature of obesity and its associated systemic complications.

### Ethical Approvals and Compliance

This study was conducted in accordance with established ethical guidelines for animal research and approved by the Institutional Animal Ethics Committee (IAEC). The study adhered to the regulations outlined by the **Committee for Control and Supervision of Experiments on Animals (CCSEA)** under Animal License No. 2173/PO/RcBiBt/S/2022/CCSEA, dated 27th June 2022. The protocol for the study was reviewed and approved by the IAEC under Protocol No.: GVSAP/IAEC/2024-035. The study followed guidelines to minimize pain, discomfort, and stress to the animals.

### Animal Welfare Measures

Animals were monitored twice daily for clinical signs. Specific interventions were planned for animals in anticipation of any distress during treatments. Humane endpoints were established to minimize suffering, including early euthanasia in cases of severe distress or body weight loss exceeding acceptable limits. All animals in the control and CCT-217 treated groups remained clinically healthy during the whole of the study and at the end, euthanasia was performed via an overdose of isoflurane, ensuring a painless and humane end. Tissue samples were collected post-mortem for further analysis. Animal housing and enrichment were designed to provide a comfortable and stimulating environment, following CCSEA and IAEC recommendations.

## Authors’ Contributions

**Raj Reddy:** Conceived the idea, provided oversight for the study, and contributed to the design of the research. Played a leading role in supervising the execution of the study, interpreting results, and editing the manuscript. **Inderpal Singh:** Designed the research, planned the study, and performed data analysis as the Data Scientist and Bioinformatician. Managed the project execution, drafted and revised the manuscript..

